# Bipartite chromatin recognition by Hop1 from two diverged holozoa

**DOI:** 10.1101/2025.06.14.659699

**Authors:** Alyssa A. Rodriguez, Alessandro E. Cirulli, Katie Chau, Justin Nguyen, Qiaozhen Ye, Kevin D. Corbett

## Abstract

In meiosis, ploidy reduction is driven by a complex series of DNA breakage and recombination events between homologous chromosomes, orchestrated by meiotic HORMA domain proteins (HORMADs). Meiotic HORMADs possess a central chromatin binding region (CBR) whose architecture varies across eukaryotic groups. Here, we determine high-resolution crystal structures of the meiotic HORMAD CBR from two diverged aquatic holozoa, *Schistosoma mansoni* and *Patiria miniata*, which reveal tightly-associated PHD and winged helix-turn-helix (wHTH) domains. We show that PHD–wHTH CBRs bind duplex DNA through their wHTH domains, and identify key residues that disrupt this interaction. Combining experimental and predicted structures, we show that the CBRs’ PHD domains likely interact with the tail of histone H3, and may discriminate between unmethylated and trimethylated H3 lysine 4. Finally, we show that holozoa Hop1 CBRs bind nucleosomes *in vitro* in a bipartite manner involving both the PHD and wHTH domains. Our data reveal how meiotic HORMADs with PHD–wHTH CBRs can bind chromatin and potentially discriminate between chromatin states to drive meiotic recombination to specific chromosomal regions.

## INTRODUCTION

Meiosis is a specialized two-stage cell division program that decreases chromosome number (ploidy) by half to produce haploid gametes or spores (Zickler & Kleckner, 2023). Ploidy reduction in meiosis requires the introduction of programmed DNA double-strand breaks (DSBs) along each chromosome, followed by the repair of these breaks as interhomolog crossovers or chiasmata. Crossovers enable the proper association and then segregation of homologous chromosomes from one another in the meiosis I division, and also drive genetic diversity in offspring (Raghavan & Hochwagen, 2025). During early prophase of meiosis I, each pair of replicated sister chromosomes condenses into a linear array of chromatin loops around a protein assembly termed the meiotic chromosome axis (Ur & Corbett, 2021; Ito & Shinohara, 2022). The chromosome axis is highly conserved across eukaryotes and typically contains three major components: HORMA domain containing proteins (HORMADs) (Gu *et al*, 2022; Prince & Martinez-Perez, 2022), axis core proteins that form filamentous assemblies through coiled-coil domains (West *et al*, 2019), and DNA-binding cohesin complexes with at least one meiosis-specific subunit (Sakuno & Hiraoka, 2022; Ur & Corbett, 2021). HORMADs play multiple roles in meiotic prophase, including recruiting DSB machinery to form DNA breaks at specific locations called DSB hotspots (Panizza *et al*, 2011; Woltering *et al*, 2000; Milano *et al*, 2024). Axis core proteins recruit HORMADs to chromatin through conserved “closure motifs” and cooperate with cohesin complexes to assemble regular chromatin loop arrays, supporting recombination and homolog pairing/synapsis (Gu *et al*, 2022; Ur & Corbett, 2021; Woltering *et al*, 2000; Kim *et al*, 2014; West *et al*, 2019).

The *S. cerevisiae* meiotic HORMAD protein Hop1 was recently found to contain a central chromatin binding region (CBR) that specifically binds nucleosomes, the fundamental unit of eukaryotic chromatin (Milano *et al*, 2024; Heldrich *et al*, 2022). The *S. cerevisiae* Hop1 CBR comprises a PHD domain (plant homeodomain) tightly packed against a variant winged helix-turn-helix domain (wHTH), plus an extended C-terminal region (HTH-C) that drapes across both domains. PHD domains typically bind histone tails, often histone H3, and can specifically recognize modifications like trimethylation at residue lysine 4 (H3K4me3) (Sanchez & Zhou, 2011). Notably, the canonical lysine binding pocket in the PHD domain of *S. cerevisiae* and closely related budding-yeast Hop1 CBRs is poorly conserved, suggesting that these proteins do not bind histone tails (Milano *et al*, 2024). Indeed, a structure of the *S. cerevisiae* Hop1 CBR bound to a nucleosome shows that Hop1 binds the outer surface of the highly bent nucleosomal DNA through a noncanonical DNA binding surface involving both the PHD and wHTH domains (Milano *et al*, 2024). The *S. cerevisiae* Hop1 CBR mediates a general enrichment of chromosome axis proteins in nucleosome-rich regions of the genome (Milano *et al*, 2024), but apparently does not recognize a specific chromatin state.

While *S. cerevisiae* and other budding yeast possess a Hop1 CBR with a PHD–wHTH–HTH-C architecture, HORMAD proteins in other eukaryotes possess CBRs with distinct architectures (Milano *et al*, 2024). While HORMADs from major animal model organisms like *M. musculus* and *C. elegans* lack the CBR entirely, HORMADs in some holozoa possess a PHD–wHTH CBR (lacking the HTH-C extension found in budding yeast). Notably, sequence alignments of PHD–wHTH CBRs from holozoa show that these proteins’ PHD domain lysine binding pockets are highly conserved, suggesting that they may bind chromatin in a manner distinct from *S. cerevisiae* Hop1, and potentially recognize particular histone modifications (Milano *et al*, 2024). Meanwhile, HORMAD proteins in archaeplastida (plants and algae) often possess a CBR with tandem wHTH domains (Milano *et al*, 2024). These findings suggest that HORMAD CBRs from different eukaryotic groups recognize chromatin through distinct mechanisms.

Here, we show by x-ray crystallography that the Hop1 CBRs from two diverged holozoa - the blood fluke *Schistostoma mansoni* and the sea star *Patiria miniata* - are structurally similar to one another, comprising tightly-packed PHD and wHTH domains. In contrast to the budding yeast Hop1 CBR, we find that *S. mansoni* and *P. miniata* Hop1 CBRs bind DNA through their wHTH domains through a canonical DNA binding interface. We further find that the *P. miniata* Hop1 CBR binds nucleosomes, and that both the PHD and wHTH domains contribute to binding, suggesting a bipartite recognition mechanism for chromatin binding. Overall, our data show that despite the architectural variability of meiotic HORMAD CBRs, they broadly share the ability to recognize and bind chromatin.

## RESULTS AND DISCUSSION

### Structure of PHD–wHTH Hop1 CBRs from two diverged holozoa

Our prior study of the budding yeast Hop1 CBR showed that this module, which comprises tightly-packed PHD, wHTH, and HTH-C domains, binds bent nucleosomal DNA in a non-canonical manner through a composite interface spanning its PHD and wHTH domains (Milano *et al*, 2024). Our phylogenetic analysis also revealed that many holozoa possess Hop1 CBRs with PHD and wHTH domains, and lack the HTH-C extension (**Figure 1A-B**) (Milano *et al*, 2024). To determine these proteins’ structures and mechanisms of chromatin binding, we expressed and purified the Hop1 CBRs of two diverged aquatic holozoa: *Schistosoma mansoni* (blood fluke) and *Patiria miniata* (star fish). We crystallized and determined high-resolution x-ray crystal structures of both proteins using single-wavelength anomalous diffraction (SAD) methods with the anomalous diffraction of endogenously-bound Zn^2+^ ions. The structures show CBRs with PHD and wHTH domains tightly packed on one another in an overall configuration similar to that of the budding yeast Hop1 CBR, but lacking the HTH-C extension seen in that protein (**Figure 1C-F, Figure S1**). The *S. mansoni* and *P. miniata* Hop1 CBRs are 43% identical at the amino acid level, and are nearly identical in structure, with an overall Cɑ root mean squared displacement (rmsd) of 1.4 Å over 136 residues. Within the CBR, the N-terminal PHD domain shows two highly conserved zinc coordination sites, one with three cysteines and one histidine (*S. mansoni* Hop1 residues C284, C286, H306, C309) and the second with four cysteines (*S. mansoni* Hop1 C298, C301, C324, C327) (**Figure 1D,F**).

**Figure 1.**
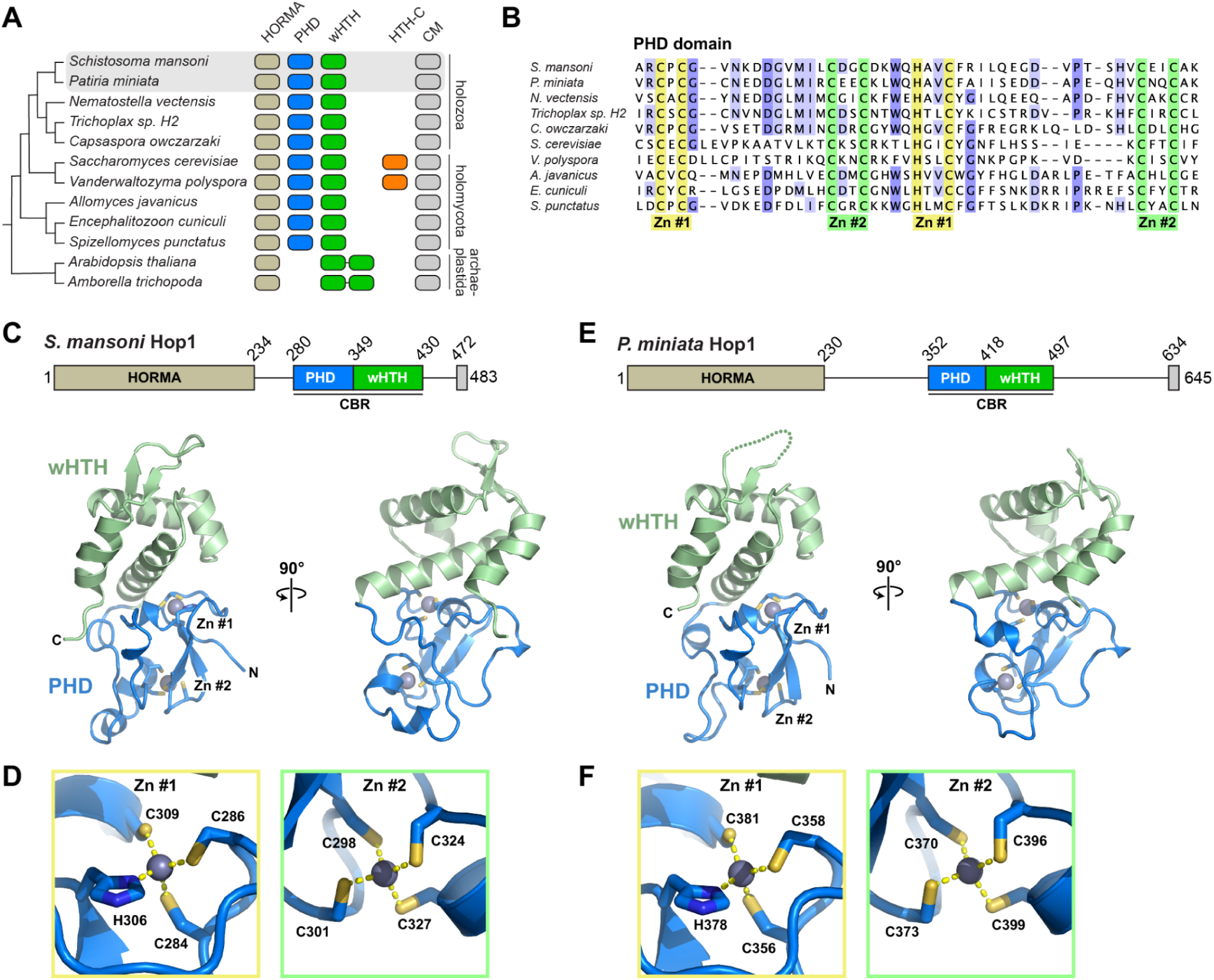
Structures of the *S. mansoni* and *P. miniata* Hop1 CBRs. **(A)** Domain architecture of Hop1 from selected eukaryotes that encode a chromatin binding region (CBR), with presence of a HORMA domain indicated in light brown, a plant homeodomain (PHD) in blue, winged helix-turn-helix (wHTH) in green (two organisms possess tandem wHTH domains and lack a PHD domain), HTH-C C-terminal chromatin binding region extension in orange, and a HORMA domain-binding closure motif (CM) in gray. **(B)** Sequence alignment of PHD domains from organisms shown in panel (A). Conserved Zn^2+^-binding residues are highlighted in yellow (Zn #1) and green (Zn #2). NCBI accession numbers for displayed sequences are XP_018653011 (*S. mansoni*), XP_038051536.1 (*P. miniata*), XP_001639673 (*N. vectensis*), RDD46648 (*Trichoplax* sp. H2), XP_004363476 (*C. owczarzaki*), NP_012193 (*S. cerevisiae*), XP_001642921 (*V. polyspora*), KAJ3363264 (*A. javanicus*), CAD25118 (*E. cuniculi*), and KND01595 (*S. punctatus*). **(C)** Crystal structure of the *Schistosoma mansoni* Hop1 CBR, with PHD domain blue, wHTH domain green, and bound Zn^2+^ ions shown as gray spheres. **(D)** Closeup view of bound Zn^2+^ ions for the *Schistosoma mansoni* Hop1 CBR, with Zn^2+^-coordinating residues shown as sticks. **(E)** Crystal structure of the *Patiria miniata* Hop1 CBR, with PHD domain blue, wHTH domain green, and bound Zn^2+^ ions shown as gray spheres. **(F)** Closeup view of bound Zn^2+^ ions for the *Patiria miniata* Hop1 CBR, with Zn^2+^-coordinating residues shown as sticks.

### The Hop1 CBR wHTH domain binds DNA

wHTH domains canonically bind DNA via insertion of an ɑ-helix (recognition helix 3) into the major groove of DNA, plus interaction of the “wing” motif with the neighboring minor groove (Lai *et al*, 1993; Gajiwala & Burley, 2000). Structure-similarity searches using DALI (Holm, 2022) and Foldseek (van Kempen *et al*, 2024) revealed that the *S. manoni* and *P. miniata* Hop1 CBR wHTH domains are more structurally similar to canonical DNA-binding wHTH domains than to the budding-yeast Hop1 CBR, which binds DNA in a non-canonical manner (Milano *et al*, 2024). We modeled DNA-bound *S. mansoni* and *P. miniata* Hop1 CBRs by overlaying their wHTH domains with a structure of DNA-bound *H. sapiens* hepatocyte nuclear factor-3 (HNF-3; PDB ID 1VTN) (Clark *et al*, 1993). The resulting models (**Figure 2A-B**) show that the Hop1 CBRs can accommodate DNA on a positively-charged surface (**Figure 2C-D**), supporting the idea that these proteins bind DNA via the canonical wHTH-DNA binding interface. To test for DNA binding, we used a fluorescence polarization-based assay with a fluorescein-labeled double-stranded DNA oligonucleotide and His-GST-tagged Hop1 CBRs. We detected DNA binding by both *S. mansoni* and *P. miniata* Hop1 CBRs (*Kd* = 16 µM for both; **Figure 2E-F**). Mutation of arginine residues in each protein’s predicted DNA binding surface (R389 and R407 for *S. mansoni*; R463, R477, and R463 for *P. miniata*) to alanine resulted in a complete loss of DNA binding (**Figure 2E-F**). Overall, these data show that, in contrast to budding-yeast Hop1 CBRs, *S. mansoni* and *P. miniata* Hop1 CBRs bind DNA via a canonical wHTH-DNA binding interface.

**Figure 2.**
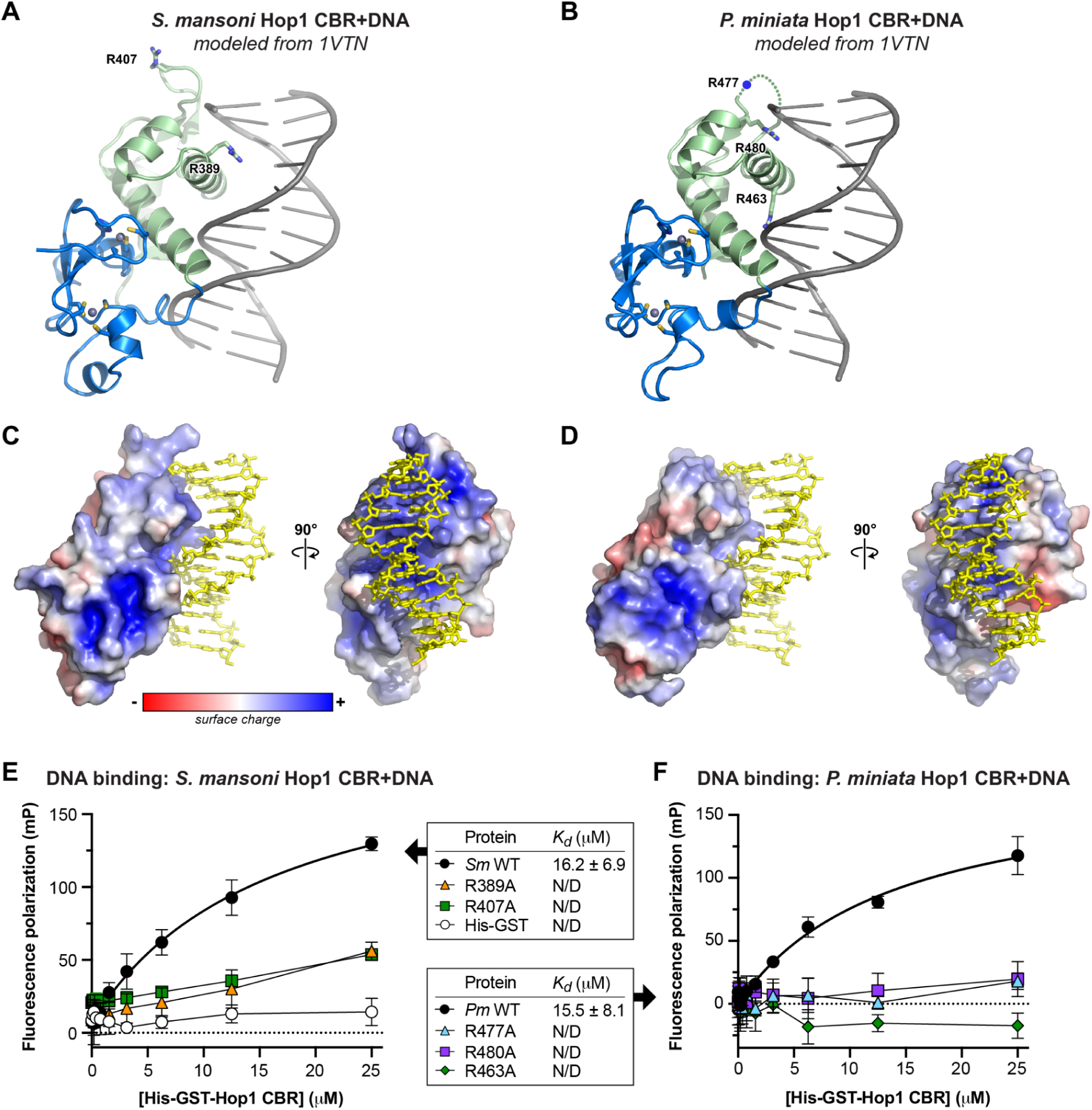
Holozoa Hop1 CBRs bind DNA through their wHTH domains. **(A)** Crystal structure of *S. mansoni* Hop1 CBR modeled with DNA (shown in gray) by overlaying with a crystal structure of a DNA-bound wHTH domain (HNF-3; PDB ID 1VTN) (Clark *et al*, 1993). Residues R389 and R407 are shown as sticks. **(B)** Crystal structure of *P. miniata* Hop1 CBR modeled with DNA as in panel (A). Residues R463 and R480 are shown as sticks; the location of R477 (in the disordered “wing”) is indicated with a blue circle. **(C)** Two views of an electrostatic surface of *S. mansoni* Hop1 CBR (calculated with APBS) (Jurrus *et al*, 2018), with modeled DNA shown in yellow sticks. **(D)** Two views of an electrostatic surface of *P. miniata* Hop1 CBR (calculated with APBS), with modeled DNA shown in yellow sticks. **(E)** Fluorescence polarization assay showing binding of His-GST-tagged *S. mansoni* Hopo1 CBR to a 20-base pair double-stranded DNA. Wild-type Hop1 is indicated in black circles, R389A in orange triangles, R407A in green squares, and His-GST negative control in white circles. **(F)** Fluorescence polarization assay showing binding of His-GST-tagged *P. miniata* Hop-1 CBR to a 20-base pair double-stranded DNA. Wild-type Hop1 is indicated in black circles, R477A in cyan triangles, R480A in purple squares, and R463A in green diamonds.

### The Hop1 CBR PHD domain likely binds a histone tail

PHD domains have diverse roles in chromatin remodeling and transcription, and typically function by binding a modified or unmodified lysine residue on a histone tail in a conserved hydrophobic pocket. PHD domains predominantly bind trimethylated histone H3 lysine 4 (H4K4me3), though some PHD domains bind unmethylated (me0), monomethylated (me1), or dimethylated (me2) H3K4 or other histone tail residues like H3 arginine 2 or lysine 14 (Sanchez & Zhou, 2011). The lysine-binding pocket of PHD domains typically comprises an “aromatic cage” with hydrophobic or aromatic residues in four conserved positions termed I, II, III, and IV (Sanchez & Zhou, 2011; Ramón-Maiques *et al*, 2007). Position I is typically a tryptophan, and position II is often a methionine, which is proposed to contribute to binding through dispersion forces and sulfur-pi interactions (Albanese & Waters, 2021). Positions III and IV are more variable, with position III mostly hydrophobic and position IV either hydrophobic or negatively-charged.

Structure-based similarity searches with DALI (Holm, 2022) and Foldseek (van Kempen *et al*, 2024) revealed that *S. mansoni* and *P. miniata* Hop1 CBR PHD domains are most closely related to a set of PHD domains that bind H3K4me3 (*S. cerevisiae* Set3 (PDB ID 5TDW) (Gatchalian *et al*, 2017), *H. sapiens* MLL5 (PDB ID 4L58) (Ali *et al*, 2013)) or H3K4me2 (*H. sapiens* PHF20 (PDB ID 5TAB) (Klein *et al*, 2016), *H. sapiens* ASH1L (PDB IF 7Y0I) (Yu *et al*, 2022)). Comparison of these proteins’ lysine binding pockets with our Hop1 CBR structures revealed strong similarity: position I is a tryptophan in all cases, position II is a methionine in most cases (threonine in Set3), position III is typically a small hydrophobic residue (alanine or valine in Hop1; isoleucine, valine, or threonine in the others), and position IV is negatively-charged (aspartate or glutamate; **Figure 3A-D**).

**Figure 3.**
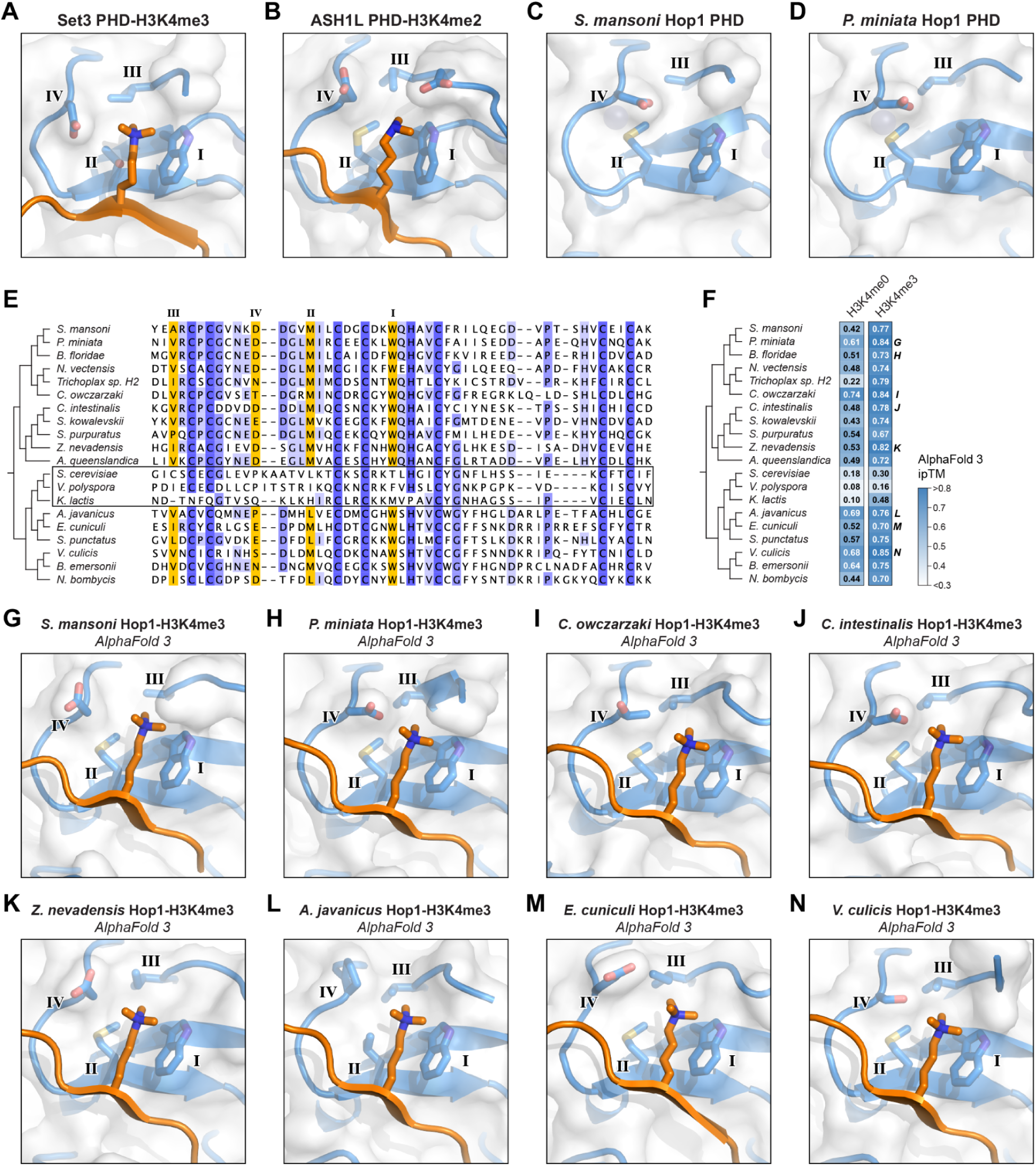
Hop1 CBRs likely bind a histone tail. **(A)** Closeup of *S. cerevisiae* Set3 PHD domain (blue) binding to trimethylated lysine 4 of histone H3 (H3K4me3; orange) (PDB ID 5TDW) (Gatchalian *et al*, 2017), with canonical PHD domain lysine binding pocket motif residues I, II, III, and IV labeled (Sanchez & Zhou, 2011). **(B)** Closeup of *H. sapiens* ASH1L PHD domain (blue) binding to H3K4me2 (orange) (PDB ID 7Y0I) (Yu *et al*, 2022), with canonical PHD domain lysine binding pocket motif residues I, II, III, and IV labeled. **(C)** Closeup of the *S. mansoni* Hop1 CBR PHD domain lysine binding pocket with canonical PHD domain lysine binding pocket motif residues I, II, III, and IV labeled. **(D)** Closeup of the *P. miniata* Hop1 CBR PHD domain lysine binding pocket with canonical PHD domain lysine binding pocket motif residues I, II, III, and IV labeled. **(E)** Sequence alignment of 20 Hop1 CBR PHD domains, with canonical PHD domain lysine binding pocket motif residues I, II, III, and IV highlighted in orange. Boxed are three Hop1 CBRs from budding yeast that show the PHD–wHTH–HTH-C architecture, for which the lysine binding pocket is not well conserved (Milano *et al*, 2024). See **Methods** for NCBI accession numbers. **(F)** Summary of AlphaFold 3 predictions of the interaction between Hop1 CBRs and the histone H3 N-terminal tail, either unmodified (H3K4me0) or with lysine 4 trimethylated (H3K4me3). Shown are ipTM scores for the most confident of five models generated; see all scores and scores for histones H2A, H2B, and H4 tails in **Figure S2A**. Models above the medium-confidence ipTM cutoff of 0.6 are indicated with white text. **(G-N)** Closeup of 8 Hop1 CBR PHD domains (blue) binding H3K4me3, as predicted by AlphaFold 3. Canonical PHD domain lysine binding pocket motif residues I, II, III, and IV labeled. See **Figure S2B-I** for the same views colored by confidence (pLDDT), and **Figure S3** for complete models and PAE plots.

We next performed a sequence alignment of a broad set of PHD domains found in eukaryotic Hop1 CBRs (Milano *et al*, 2024). This alignment shows high conservation in positions I-IV among Hop1 CBRs from holozoa and holomycota, with the exception of the budding-yeast family whose CBRs possess the HTH-C extension (**Figure 3E**). In Hop1 CBRs with the PHD–wHTH architecture, position I is highly conserved as a tryptophan, position II is a methionine or leucine, position III is a small hydrophobic residue (alanine, valine, or isoleucine), and position IV is most often an aspartate. These data show that, with the exception of budding-yeast Hop1 CBRs which are known to bind nucleosomes in a non-canonical manner (Milano *et al*, 2024), the PHD domain lysine binding pockets in Hop1 CBRs are highly conserved and similar to PHD domains known to bind di- or trimethylated H3K4.

We attempted to directly detect binding between purified *S. mansoni* or *P. miniata* Hop1 CBRs and diverse modified and unmodified histone tail peptides, without success. As an alternative, we used AlphaFold 3 (Abramson *et al*, 2024) to predict the structures of 20 different Hop1 CBRs with the N-terminal tails of histones H2A, H2B, H3 (unmodified or trimethylated at lysine 4), and H4, then systematically analyzed the confidence of the predicted interaction using the AlphaFold ipTM (interface predicted template modeling) score (**Figure 3F, S2A**). Our test set included 17 Hop1 CBRs with PHD and wHTH domains (including *S. mansoni* and *P. miniata*), and three budding-yeast Hop1 CBRs with PHD, wHTH, and HTH-C domains. Since PHD–wHTH–HTH-C CBRs bind nucleosomes in a non-canonical manner (Milano *et al*, 2024) and do not possess a conserved lysine binding pocket in their PHD domains (**Figure 3E**), these proteins served as a negative control. Across all 17 Hop1 CBRs with PHD–wHTH architecture, the most confidently-predicted histone tail interactions were with histone H3 with trimethylated lysine 4 (H3K4me3). All 17 predictions showed an ipTM score above the medium-confidence threshold of 0.6, and four scored above the high-confidence threshold of 0.8 (**Figure 3F**) (Kim *et al*, 2024). All 17 predictions showed the modified H3K4me3 residue docked in the CBR’s lysine binding pocket, consistent with known structures of PHD-H3K4me3 complexes (**Figure 3G-N, S2B-I, S3**). Meanwhile, the three Hop1 CBRs from budding yeast consistently showed low-confidence predictions with all histone tails, including the H3 tail with H3K4me3 (**Figure 3F, S2A**). Overall, these predictions are consistent with a model in which PHD–wHTH CBRs can bind a histone tail residue, potentially trimethylated H3K4, as part of its chromatin recognition mechanism.

### Hop1 CBRs from holozoa bind nucleosomes

We previously showed that the budding yeast Hop1 PHD–wHTH–HTH-C CBR binds nucleosomes via a non-canonical surface, recognizing the bent nucleosomal DNA and not contacting the histone proteins themselves (Milano *et al*, 2024). To test if the holozoa Hop1 PHD–wHTH CBR domain binds nucleosomes, we performed electrophoretic mobility shift assays (EMSAs) with recombinant *P. miniata* Hop1 CBR and nucleosome core particles (NCPs) assembled from *Xenopus laevis* histones and DNA containing the WIdom 601 nucleosome-positioning sequence, either unmodified or containing a structural analog of the H3K4me3 modification (Simon *et al*, 2007). We found that the *P. miniata* Hop1 CBR binds nucleosomes with an affinity in the low micromolar range, and that alanine mutants in the PHD domain aromatic cage (position I residue W376) and the wHTH domain DNA binding surface (R480) each show reduced nucleosome binding (**Figure 4A-B**). Wild-type *P. miniata* Hop1 CBR bound to unmodified and H3K4me3 nucleosomes with roughly equivalent affinity (**Figure 4C**), indicating the Hop1 CBR alone does not strongly discriminate between H3K4 methylation states *in vitro*. We also performed EMSAs with the isolated Widom 601 DNA, and observed that the wHTH domain mutant R480A, but not the PHD domain mutant W376A, reduced DNA binding (**Figure 4D**). Overall, these data are consistent with bipartite recognition of nucleosomes by the *P. miniata* Hop1 CBR, with both the PHD and wHTH domains contributing to the interaction.

**Figure 4.**
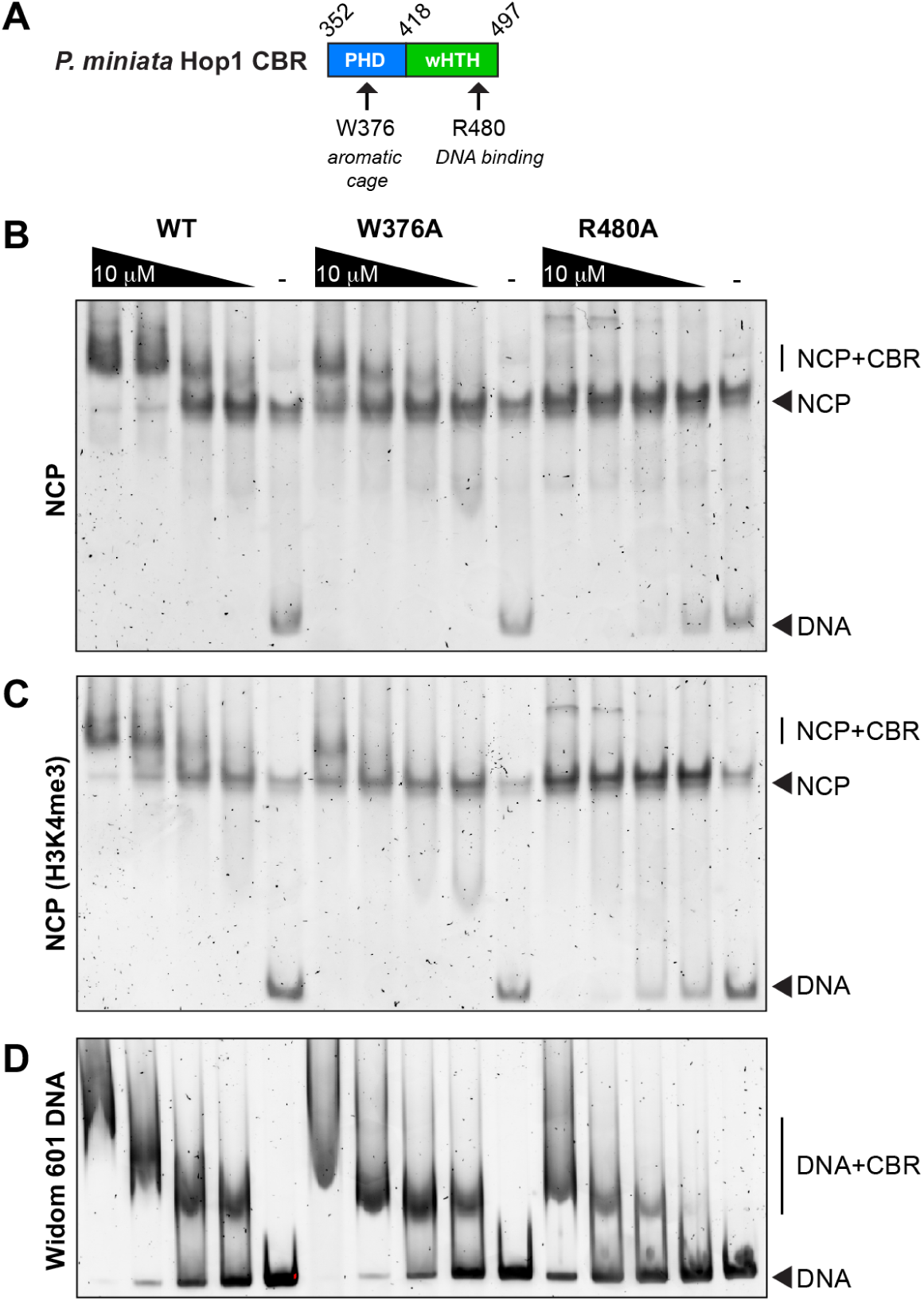
Bipartite nucleosome recognition by the *P. miniata* Hop1 CBR. **(A)** Schematic of the *P. miniata* Hop1 CBR and residues mutated in the PHD domain aromatic cage (W376) and wHTH domain DNA binding surface (R480). **(B)** Representative electrophoretic mobility shift assay (EMSA) with *P. miniata* Hop1 CBR (Wild type (WT), W376A, or R480A at 10 μM and two-fold dilutions) and 50 nM nucleosome core particles (NCPs). **(C)** Representative EMSA with *P. miniata* Hop1 CBR and NCPs containing a H3K4me3 modification analog. **(D)** Representative EMSA with *P. miniata* Hop1 CBR and Widom 601 DNA.

In conclusion, we show that meiotic HORMAD CBRs with the PHD–wHTH architecture recognize chromatin through a bipartite interface, with histone tail binding by the PHD domain and DNA binding by the wHTH domain. While our *in vitro* assays do not show strong discrimination between methylated and unmethylated H3K4 by the Hop1 CBR, our AlphaFold predictions nonetheless suggest that *in vivo*, this domain may preferentially bind chromatin with the H3K4me3 mark. Given that meiotic HORMADs promote DSB formation by recruiting DSB machinery, HORMAD CBRs with the PHD–wHTH architecture may therefore represent one mechanism for targeting DSBs to H3K4me3 chromatin. Testing this idea in a genetically-tractable model organism that possesses a PHD–wHTH CBR, such as the sea star *P. miniata* (Meyer and Hinman 2022; Zueva and Hinman 2023), is therefore an important avenue for future work. Relatedly, whether and how wHTH–wHTH CBRs found in meiotic HORMADS in archaeplastida recognize particular chromatin states will also be important to test.

## METHODS

### Cloning, expression, and protein purification

Codon-optimized gene blocks encoding the CBRs of *S. mansoni* Hop1 (NCBI XP_018653011, residues 279-430) and *P. miniata* Hop1 (NCBI XP_038051536.1, residues 352-499) were purchased from GenScript and cloned into UC Berkeley Macrolab vector 2B-T (Addgene #29666), which encodes an N-terminal TEV protease-cleavable His_6_-tag; 2C-T (Addgene #29706), which encodes an N-terminal TEV protease-cleavable His_6_-maltose binding protein (MBP) tag; and 2G-T (Addgene #29707), which encodes an N-terminal TEV protease-cleavage His_6_-glutathione S-transferase (GST) tag. Plasmids were transformed into Rosetta 2 (DE3) pLysS *E. coli* competent cells (EMD Millipore), and 5 mL Luria broth (LB) cultures were grown at 37°C for 16 hours with appropriate antibiotics. Overnight cultures were used to inoculate 1 L cultures in 2XYT media and grown at 37°C with shaking at 180 RPM until OD_600_= 0.5. Protein expression was induced with the addition of 0.2 mM Isopropyl β-d-1-thiogalactopyranoside (IPTG), the temperature shifted to 20°C, and cultures were grown an additional 16 hours. Cells were harvested by centrifugation and resuspended in lysis buffer (20 mM HEPES-NaOH pH 7.0, 300 mM NaCl, 10% glycerol, 5 mM imidazole, 10 µM ZnCl_2_).

Resuspended cells were lysed by sonication (Branson Sonifier), then the cell lysate was clarified by centrifugation. The supernatant was loaded onto an Ni^2+^ affinity column (Qiagen Ni-NTA Superflow) in lysis buffer, washed with wash buffer (20 mM HEPES-NaOH pH 7, 25 mM imidazole, 300 mM NaCl, 5 mM MgCl_2_, 10 µM ZnCl_2_, 10% glycerol, and 5 mM β-mercaptoethanol), then eluted in elution buffer (20 mM HEPES-NaOH pH 7, 500 mM imidazole, 300 mM NaCl, 5 mM MgCl_2_, 10 µM ZnCl_2_, 10% glycerol, 5 mM β-mercaptoethanol. Fractions were pooled and diluted to 100 mM NaCl through the addition of dilution buffer (20 mM HEPES-NaOH pH 7, 25 mM imidazole, 5 mM MgCl_2_, 10 µM ZnCl_2_, 10% glycerol, 5 mM β-mercaptoethanol) before loading onto a cation-exchange column (Hi-Trap SP, Cytiva #17-1152-01. Protein was eluted with a linear gradient from 100 mM to 1 M NaCl. For crystallography, His_6_-tagged protein was incubated with TEV protease (expressed and purified in-house from vector pRK793, Addgene #8827) (Kapust *et al*, 2001) for 48 hours at 4°C. For biochemical assays, His_6_-GST-tagged protein was not cleaved with TEV protease. TEV cleavage reactions were passed over a Ni^2+^ affinity column a second time to remove His_6_-tagged TEV protease and uncleaved Hop1, and the flow-through was collected and concentrated. Finally, proteins were passed over a size exclusion column (Superdex 200 Increase 10/300 GL, Cytiva). Fractions were pooled and concentrated, then stored at -80°C (for biochemical assays) or buffer-exchanged into crystallography buffer (20 mM HEPES-NaOH pH 7, 200 mM NaCl, 5 mM MgCl_2_, and 1 mM Tris-(2-Carboxyethyl)phosphine (TCEP).

### X-ray crystallography

For crystallization of *S. mansoni* Hop1 CBR, purified protein at 10 mg/mL was mixed 1:1 with well solution containing 0.1 M MES pH 6.0, 0.1 M sodium acetate, and 25% PEG 3350 in hanging drop format at 25°C. For crystallization of *P. miniata* Hop1 CBR, protein was subjected to reductive lysine methylation (Kim *et al*, 2008), then protein at 10 mg/mL was mixed 1:1 with well solution containing 0.1 M Bis Tris pH 6.5, 0.1 M potassium thiocyanate, and 17% PEG 3350 in hanging drop format at 25°C. In both cases, crystals were cryoprotected with an additional 30% xylitol and flash-frozen in liquid nitrogen. Diffraction data were collected at Advanced Photon Source beamline 24ID-C and processed with the RAPD pipeline, which uses XDS (Kabsch, 2010) for indexing and integration, AIMLESS (Evans & Murshudov, 2013) for scaling, and TRUNCATE for conversion to structure factors (Agirre *et al*, 2023). Anomalous sites representing protein-bound zinc ions were identified with hkl2map/SHELX (Sheldrick, 2010) and input into the Phenix Autosol pipeline (Liebschner *et al*, 2019) for phasing and automatic model building. Initial models were manually rebuilt in COOT (Emsley *et al*, 2010) and refined in phenix.refine (Adams *et al*, 2010). *S. mansoni* Hop1 CBR (1.45 Å resolution) was refined using positional and individual anisotropic B-factor refinement. *P. miniata* Hop1 CBR (1.84 Å resolution) was refined using positional, individual isotropic B-factor, and TLS refinement (one TLS group per protein chain) (**Table 1**). Both models were refined with riding hydrogen atoms. All structural figures were created using PyMol (version 3; Schrödinger, LLC). Surface charge representations were calculated with APBS (Jurrus *et al*, 2018).

**Table 1.** Crystallographic data and refinement.

|  | *S. masoni* Hop1 CBR | *P. miniata* Hop1 CBR |
| --- | --- | --- |
| **Data Collection** |  |  |
| Synchrotron/Beamline | APS-24ID-C | APS-24ID-C |
| Date collected | 9/22/2021 | 9/22/2021 |
| Resolution (Å) | 64.96 - 1.45 | 82.95-1.84 |
| Wavelength (Å) | 1.2827 | 1.2827 |
| Space Group | P2<sub>1</sub>2<sub>1</sub>2<sub>1</sub> | P2<sub>1</sub>2<sub>1</sub>2<sub>1</sub> |
| Unit Cell Dimensions (a,b,c) Å | 33.03, 64.96, 65.26 | 38.65, 106.96, 131.40 |
| Unit cell Angles ($\alpha,\beta,\gamma$)° | 90.00, 90.00, 90.00 | 90.00, 90.00, 90.00 |
| I/$\sigma$ (last shell) | 30.9 (5.3) | 14.7 (1.3) |
| R<sub>merge</sub> (last shell) | 0.08 (0.458) | 0.067 (0.827) |
| R<sub>meas</sub> (last shell) | 0.088 (0.573) | 0.080 (1.018) |
| CC<sub>1/2</sub> (last shell) | 0.997 (0.557) | 0.997 (0.454) |
| Completeness (last shell) % | 94.4 (50.3) | 98.9 (99.1) |
| Anomalous completeness (last shell) % | 90.3 (29.8) | 90.3 (75.4) |
| Number of reflections | 135024 | 153221 |
| unique | 24095 | 47903 |
| Multiplicity (last shell) | 5.6 (2.3) | 3.2 (2.7) |
| **Refinement** |  |  |
| Resolution (Å) | 1.45 | 1.84 |
| No. of reflections | 24041 (1573) | 47834 (4585) |
| working | 22883 (1475) | 45414 (4366) |
| free | 1158 (98) | 2420 (219) |
| R<sub>work</sub> (last shell) (%) | 12.88 (20.83) | 19.79 (33.89) |
| R<sub>free</sub> (last shell) (%) | 15.83 (28.34) | 23.54 (35.61) |
| **Structure/Stereochemistry** |  |  |
| No. of atoms | 1400 | 7970 |
| solvent | 216 | 326 |
| hydrogens | 1186 | 3780 |
| r.m.s.d. bond lengths (Å) | 0.014 | 0.0085 |
| r.m.s.d. bond angles (°) | 1.39 | 0.97 |
| SBGrid Data Bank ID | 1153 | 1156 |
| Protein Data Bank ID | 9MUG | 9MYP |

### AlphaFold Structure Prediction

Structure predictions for Hop1 CBR-histone tail complexes were generated using the publicly-available AlphaFold 3 server at https://alphafoldserver.com (Abramson *et al*, 2024). Template settings were set with the default PDB cut-off date of September 29, 2021, unless noted otherwise. ipTM scores for each model were extracted from the fold_job-name_summary_confidences_model-number.json files. NCBI/Uniprot accession numbers for proteins used were: XP_018653011 (*S. mansoni*), XP_038051536.1 (*P. miniata*), XP_001639673 (*N. vectensis*), XP_035662583 (*B. floridae*), XP_001639673 (*N. vectensis*), RDD46648 (*Trichoplax* sp. H2), XP_004363476 (*C. owczarzaki*), XP_009860628 (*C. intestinalis*), XP_002731279 (*S. kowalevskii*), XP_030838487 (*S. purpuratus*), A0A067R097 (Uniprot; *Z. nevadensis*), A0A1X7V (Uniprot; *A. queenslandica*), NP_012193 (*S. cerevisiae*), XP_001642921 (*V. polyspora*), XP_452539 (*K. lactis*), KAJ3363264 (*A. javanicus*), CAD25118 (*E. cuniculi*), KND01595 (*S. punctatus*), ELA45904 (*V. culicis*), KAI9179672.1 (*B. emersonii*), and EOB12410 (*N. bombycis*). Histone tail sequences used were SGRGKQGGKTRAKAKTRSSR (H2A), AKSAPAPKKGSKKAVTKTQK (H2B), ARTKQTARKSTGGKAPRKQL (H3), ART(K + N-trimethyllysine)QTARKSTGGKAPRKQL (H3K4me3), and SGRGKGGKGLGKGGAKRHRK (H4).

### Fluorescence polarization assays

To measure DNA binding, His_6_-GST-tagged *S. mansoni* and *P. miniata* Hop1 CBR at the indicated concentrations were incubated with 50 nM fluorescein-labeled DNA duplex (5’-6-FAM-CTTATATCTGAATAGTCAGT-3’ annealed with 5’-ACTGACTATTCAGATATAAG-3’) in binding buffer (20 mM Tris-HCl pH 8, 100 mM sodium glutamate, 5 mM MgCl_2_, 3% glycerol, and 0.5 mM β-mercaptoethanol). Samples were incubated at 4°C for 30 minutes before reading fluorescence polarization on a Tecan Spark plate reader. Data were analyzed with GraphPad Prism version 10 (GraphPad software) using a one-site specific binding model.

### Reconstitution of nucleosome core particles

Nucleosome core particles were reconstituted following published protocols (Luger *et al*, 1999). Briefly, lyophilized *Xenopus laevis* histones H2A, H2B, H3 (or H3K4me3), and H4 were purchased from the Histone Source at Colorado State University (https://histonesource-colostate.nbsstore.net/). The H3K4me3 histone contained K4C and C110A point mutants, and cysteine 4 was modified with a trimethyl-aminoethyl group according to published procedures (Simon *et al*, 2007) to mimic trimethyllysine. Histones were unfolded with incubation and shaking for 1 hour at room temperature in unfolding buffer (6 M guanidine hydrochloride, 20 mM Tris-HCl pH 7.5, 5 mM DTT). Histones were then added in equimolar ratio for 1 mg/mL final concentration. Histones were refolded into octamer and dialyzed in refolding buffer (2 M NaCl, 10 mM Tris-HCl pH 7.5, 1 mM EDTA, and 5 mM β-mercaptoethanol). After refolding, histones were concentrated via centrifugation and loaded onto a size exclusion column (Superdex 200 16/600) equilibrated in refolding buffer. Fractions were analyzed by SDS-PAGE, and fractions containing pure histones were pooled. For nucleosome reconstitution, the Widom 601 DNA sequence was amplified by PCR, purified, and concentrated. DNA was added to histone octamer in 1:1.2 molar ratio and dialyzed in 1.4 M KCl, 10 mM Tris-HCl pH 7.5, 0.1 mM EDTA, 1 mM DTT for 1 hour at 4°C. Low salt buffer (10 mM KCl, 10 mM Tris-HCl pH 7.5, 0.1 mM EDTA, 1 mM DTT) was slowly pumped into the high salt buffer for a few hours and then replaced with low salt buffer and dialyzed overnight. Nucleosomes were concentrated with a centrifugal filter and injected onto a size exclusion column (Superose 6, 10/300 GL) in 20 mM HEPES pH 7.5, 20 mM NaCl, 0.5 mM EDTA, 1 mM DTT. Fractions were run on (pre-run at 0.5X TBE for 150 V for 1 hour at 4°C) a 6% acrylamide TBE gel for 1 hour at 150V at 4°C. The gel was stained in SYBR Gold (1:10000) for 30 minutes while shaking in the dark. The gel was imaged with the ChemiDoc system, and pure nucleosomes were pooled and concentrated with a centrifugal filter.

### Electrophoretic mobility shift assays

Recombinantly purified His_6_-MBP-Hop1 CBR and reconstituted non-methylated and methylated (H3K4me3) nucleosome core particles were incubated for 60 minutes at 4°C in binding buffer (20 mM HEPES pH 7, 100 mM NaCl, 100 µM ZnCl_2_, 10% glycerol, 2 mM betamercaptoethanol). Samples were loaded onto 6% TB gel pH 9.3 (gel was prerun at 20 mA for 1 hour at 4°C) and run for 110 minutes at 20 mA at 4°C in running buffer (0.5X TB, pH 9.3). Gel was then incubated with shaking in SYPR Gold (Invitrogen S11494) for 10 minutes in the dark. Gel was imaged on a ChemiDOC Imaging system (ChemiDoc 12003153).

### Data Availability

Raw diffraction data have been deposited at the SBGrid Data Bank (https://data.sbgrid.org) under accession cores 1153 (*S. mansoni* Hop1 CBR) and 1156 (*P. miniata* Hop1 CBR). Reduced data and final refined crystallographic models have been deposited at the RCSB Protein Data Bank (https://rcsb.org) under accession cores 9MUG (*S. mansoni* Hop1 CBR) and 9MYP (*P. miniata* Hop1 CBR).

---

**Figure S1.**
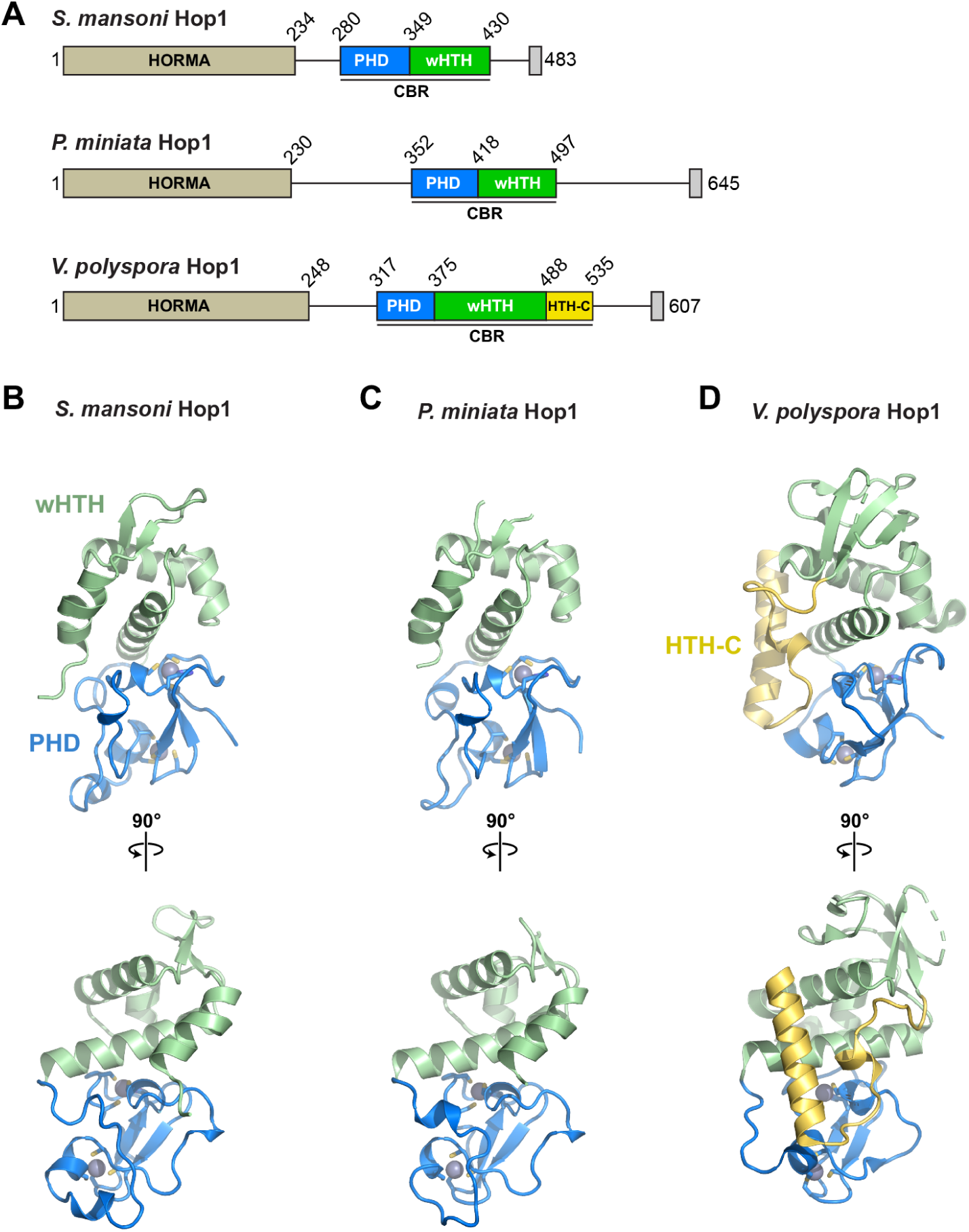
Structure of Hop1 CBRs. **(A)** Domain schematics of Hop1 CBRs from *S. mansoni*, *P. miniata*, and the budding yeast *Vanderwaltozyma polyspora*. **(B-D)** Two views of the *S. mansoni* (panel B), *P. miniata* (panel C), and *V. polyspora* (panel D; PDB ID 7UBA) (Milano *et al*, 2024), with PHD domain in blue with bound zinc ions in gray, wHTH domain in green, and HTH-C extension (only in *V. polyspora*) in yellow.

**Figure S2.**
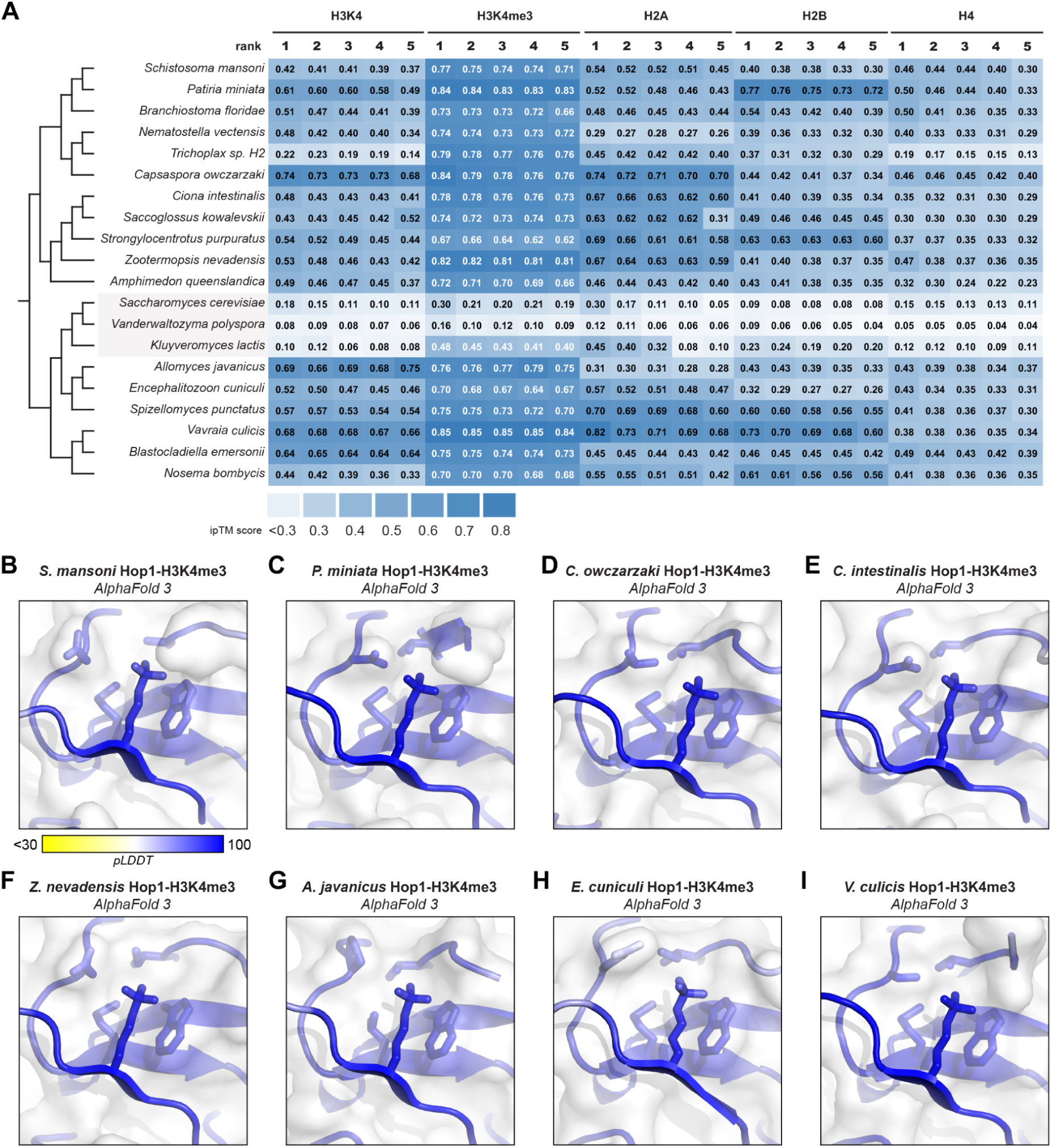
AlphaFold predictions of Hop1 CBR-histone tail complexes. **(A)** AlphaFold 3 ipTM scores for all five models generated for predicted complexes of 20 Hop1 CBRs with (from left to right) the N-terminal tails of unmodified histone H3, histone H3 with trimethylated lysine 4, histone H2A, histone H2B, and histone H4. **(B-I)** Closeup views of CBR-H3K4me3 complexes equivalent to Figure 3G-N, colored according to confidence (pLDDT: predicted local distance difference test).

**Figure S3.**
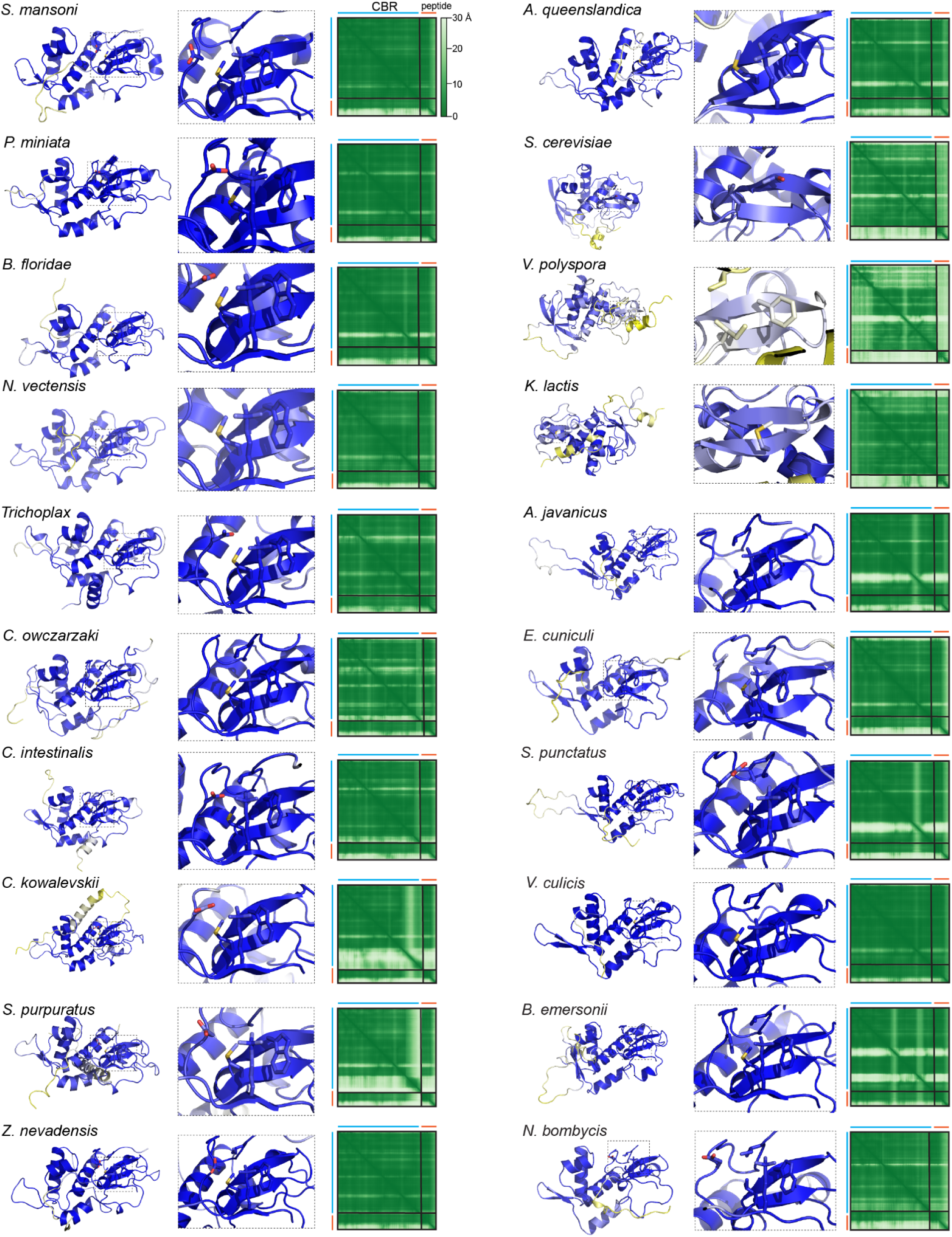
AlphaFold models of CBR-H3K4me3 tails. For each of 20 Hop1 CBR plus H3K4me3 complexes, shown are (from left to right) the overall predicted structure, a closeup of the PHD domain lysine binding pocket, and the AlphaFold 3 PAE (predicted aligned error) plot. Structures are colored according to confidence (pLDDT: predicted local distance difference test).

## ACKNOWLEDGEMENTS

The authors thank members of the Corbett lab for helpful discussions. The authors acknowledge support from the National Institutes of Health (K12 GM068524 to AAR; R35 GM144121 to KDC). This work is based upon research conducted at the Northeastern Collaborative Access Team beamlines, which are funded by the National Institute of General Medical Sciences from the National Institutes of Health (P30 GM124165). This research used resources of the Advanced Photon Source, a U.S. Department of Energy (DOE) Office of Science User Facility operated for the DOE Office of Science by Argonne National Laboratory under Contract No. DE-AC02-06CH11357.

